# LigandForge: A Web Server for Structure-Guided De Novo Drug Design

**DOI:** 10.64898/2026.03.31.715741

**Authors:** Hossam Nada, Levente Sipos-Szabó, Dávid Bajusz, György M. Keserű, Moustafa Gabr

## Abstract

Despite advances in computational drug discovery, de novo drug design remains hindered by high licensing costs and the need for specialized programming expertise. We present LigandForge, a webserver for structure-guided de novo ligand generation. LigandForge integrates structural validation and binding-site characterization; voxel-based property grid construction for spatial mapping of electrostatics and hydrophobicity; chemistry-aware fragment assembly; multi-objective lead optimization; and retrosynthetic feasibility analysis. The platform utilizes a structure-guided framework to assemble molecules from curated fragment libraries while enforcing physicochemical constraints, including molecular weight, LogP, and hybridization states. Generated molecules are refined via reinforcement learning and genetic algorithms which are subsequently evaluated using composite metrics such as the quantitative estimate of drug-likeness. By leveraging RDKit for cheminformatics and NGL viewer for real-time 3D visualization, LigandForge provides a synthesis-aware environment that bridges the gap between macromolecular structural data and experimentally feasible lead compounds without requiring local software installation.

## Introduction

Despite the significant advances in computational drug discovery, the *de novo* design of small molecules capable of targeting a defined binding site remains a challenging concept^1,2^. Targeted de novo ligand generation requires the integration of binding-site geometry, physicochemical property mapping, construction of a viable chemical starting point, and multi-objective lead optimization within a single, accessible computational environment^3,4^. While several existing frameworks and software packages address de novo drug design, their broader adoption remains limited due to significant barriers, including the requirement for specialized cheminformatics knowledge or advanced programming expertise. Another barrier toward de novo drug design is that several of the currently established commercial solutions require costly licensing agreements that render them financially less accessible to many academic institutions.

To address these limitations, we present LigandForge, an integrated, web-accessible platform designed for automated, structure-guided ligand discovery through a modular and extensible pipeline architecture (Figure 1). LigandForge framework encompasses five interconnected modules: (i) structural validation and binding-site characterization, (ii) high-dimensional voxel-based property grid construction, (iii) chemistry-aware fragment assembly, (iv) multi-objective lead optimization, and (v) retrosynthetic feasibility analysis. The server accepts protein information which is processed via a Biopython^5^ which enforces structural integrity prior to downstream analysis. Binding environments are subsequently quantified into a three-dimensional property grid through a voxel-based analyzer which enables the spatial mapping of electrostatic potentials, hydrophobic fields, and steric accessibility. These analyses are further enhanced by a custom analyzer capable of identifying hotspots and quantifying the enthalpic and entropic contribution of crystallographic water sites, providing thermodynamically grounded guidance for ligand positioning.

**Figure 1.**
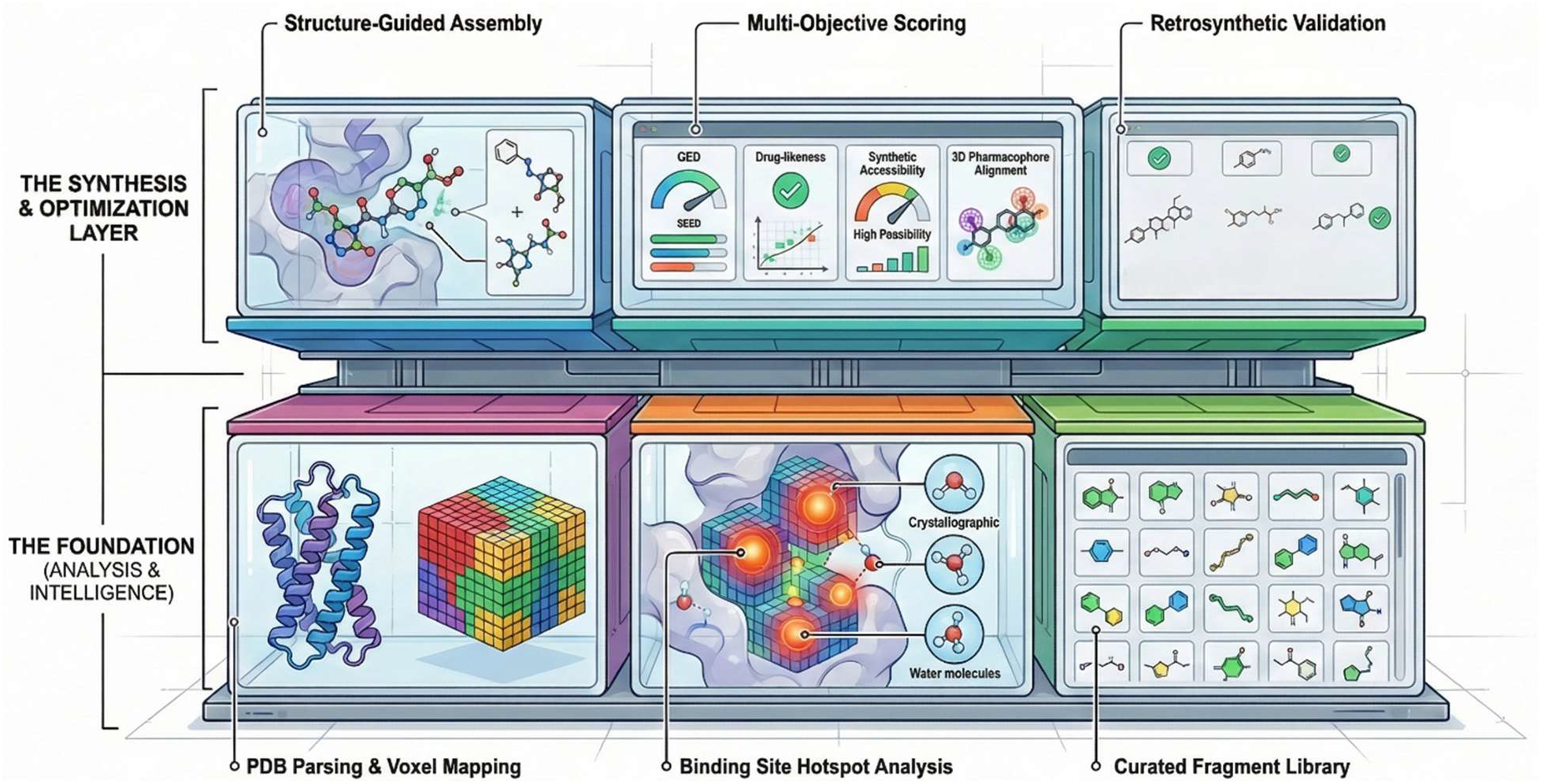
LigandForge system architecture. A schematic representation of the pipeline for structure-guided de novo design.

Small molecule generation within LigandForge is driven by a structure-guided assembly engine that performs chemistry-aware fragment attachment from a curated fragment library which is organized by functional role as core scaffolds, linkers, substituents, and bioisosteres. An attachment point manager governs growth-site selection by evaluating local hybridization states (sp, sp^2^, sp^3^), aromaticity, and available valence, while a validation engine enforces drug-likeness constraints throughout the assembly process, including molecular weight boundaries (150–700 Da), LogP ranges (−4 to 7), and rotatable bond limits. Following assembly, candidate molecules are evaluated and refined by a multi-objective scorer that computes composite molecular score metrics encompassing pharmacophore alignment, synthetic accessibility, drug-likeness as measured by the quantitative estimate of drug-likeness (QED), and structural novelty. Optimization is supported by reinforcement learning (RL), genetic algorithms (GA), and hybrid heuristic strategies, with population diversity maintained through DBSCAN clustering^6,7^ and molecular fingerprinting via the diversity manager. To ensure that computational predictions are grounded in synthetic accessibility, top-scoring candidates are subjected to retrosynthetic decomposition by a retrosynthetic analyzer, which deconstructs molecules into feasible synthetic routes and provides quantitative assessments of overall synthetic difficulty, step yield, and route feasibility.

LigandForge is implemented as a web application which is hosted on the Render server (https://ligandforge.onrender.com), enabling it to operate as a standalone web-based demonstration platform without requiring local software installation or command-line proficiency. The platform leverages RDKit for all cheminformatics computations and integrates an interactive NGL viewer for real-time three-dimensional structural validation, enabling researchers to inspect ligand–receptor complementarity directly within the browser environment. Collectively, LigandForge offers a platform for structure-guided drug discovery via a user-friendly interface that does not require programming expertise.

### Application

#### Prospective validation by co-folding

In an effort to validate its performance, we evaluated the binding affinity of LigandForge-generated molecules using Boltz-2, a state-of-the-art macromolecular structure prediction framework that includes an affinity prediction module and has been reported to achieve accuracy comparable to free energy perturbation-based methods^8^. We assessed LigandForge on three biologically and therapeutically relevant targets. Dopamine receptor D2 and epidermal growth factor receptor (EGFR), representing a GPCR and a kinase, respectively, were selected as examples of two major and commonly targeted protein classes in key therapeutic areas: neurology and oncology. We also included a more challenging target, the N-terminal domain of STAT5b, which mediates STAT5b protein oligomerization and is an emerging therapeutic target in several forms of leukemia and other cancers^9,10^. Figure 2 shows the distribution of Boltz-2-predicted affinities of all candidates from ten independent de novo generation runs (100 candidates per target), together with the predicted binding poses of molecules selected from the top-affinity candidates for each examined target.

**Figure 2.**
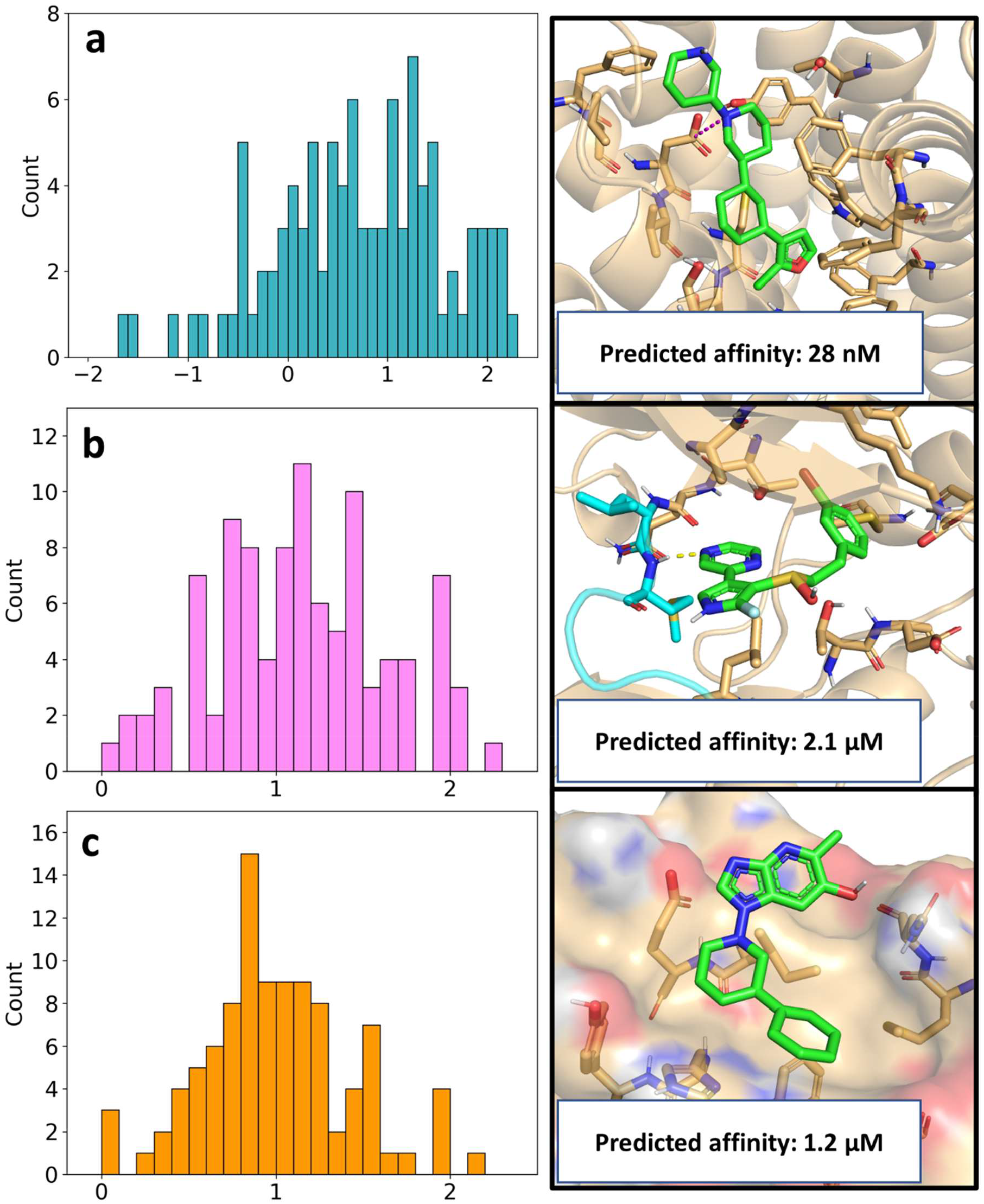
Distribution of Boltz-2 predicted affinities (logarithm of values in µM units) shown on histograms (left side), and Boltz-2 predicted binding modes of top candidates (right side) for the D2 receptor (a), EGFR kinase (b) and STAT5B N-terminal domain (c). For D2R, the key salt bridge interaction formed between D114 (D^3.32^) and the generated ligand is shown with a purple dashed line. For EGFR, the hinge region is shown in cyan, hydrogen bond interaction with the generated ligand is shown with a yellow dashed line.

LigandForge-generated molecules achieved predicted affinities in the micromolar range across all three targets, with submicromolar affinities observed for D2R. The predicted binding poses also recapitulated characteristic interaction patterns associated with the respective targets. In the case of D2R, this included the canonical salt-bridge interaction between the ligand’s tertiary amine and D144 (D^3.32^ in generic GPCR numbering)^11^. For EGFR, the generated molecules occupied the ATP-binding site and formed H-bond interactions with the hinge region, consistent with well-known binding modes of type I kinase inhibitors^12^, as well as the binding mode of erlotinib in the applied crystal structure 1M17. Notably, every generated molecule has shown a similarity of less than 0.3 to the most similar known active ligand for the corresponding target (see Supplementary Figure S1 for the three best candidates in Figure 2, as well as Supplementary Table S1 for all generated compounds). This indicates that LigandForge can generate novel molecules while still retaining favorable predicted binding properties.

## Methods

LigandForge is built on a modular, automated pipeline which is designed for structure-guided *de novo* molecular design. The pipeline integrates protein structural analysis, fragment-based assembly, and multi-objective optimization into a unified computational workflow, whose modules are summarized below.

### Target Preparation and Voxel-Based Field Analysis

Protein structures are processed by a dedicated PDB-parser, which extracts high-fidelity atom record data for protein chains, crystallographic waters, and co-crystallized ligands. A voxel-based analyzer then performs grid-based calculations within a user-defined radius of the binding site, generating a three-dimensional property grid that serves as a discrete representation of the binding pocket’s chemical environment. The property grid is initialized as five primary NumPy arrays of shape (N_x_, N_γ_, N_φ_), encoding distinct physicochemical fields. The electrostatic potential field, Φ(v) = Σ(q_i_ / ‖v − r_i_‖), is computed by summing partial charges q_i_ over all atoms at positions r_i_, weighted by inverse Euclidean distance; electrostatic variance across non-occupied voxels quantifies polar environment heterogeneity.

Lipophilic regions are mapped via hydrophobic patch identification, with the hydrophobic moment vector H_moment_ = Σ(h_v_ × (v − v_henter_)) characterizing amphipathic asymmetry within the cavity. Excluded volumes defined by protein van der Waals radii delineate physical boundaries for ligand placement, while solvent-accessible surface area and voxel-level openness (scalar 0.0–1.0) define cavity volume boundaries. Local geometric topology is encoded via shape index and curvature, derived from principal curvatures (k_1_, k_2_) of the iso surface, yielding descriptors for pocket depth, width, convexity, and distance to the nearest van der Waals surface.

The LigandForge analyzer identifies interaction hotspots in order to prioritize the binding regions which are scored on the basis of residue conservation, accessibility, and pharmacophore feature vectors. Crystallographic waters are evaluated via the water site data structure, which assigns a replaceability score and entropy contribution to each site; high-energy water positions are flagged as displacement targets, as their replacement by ligand functionality can make a substantial contribution to binding affinity^13,14^. A field complexity score derived from the Shannon entropy of the property distribution across the grid quantifies the diversity of interaction types required for complementary binding. Local field gradient vectors [dx, dy, dz] represent the direction of maximum property change and are subsequently used to orient fragments during molecular assembly.

#### Fragment-Based Molecular Assembly

Molecular generation is executed through a structure-guided assembly engine. A curated fragment library provides fragments that are categorized as core, linker, substituent, or bioisostere, and the structure guided assembly framework manages ligand growth at specific coordinates identified by the molecular elaboration engine. For each growth site at three-dimensional position P, the corresponding grid index is computed as I = ⌊(P − grid_origin) / grid_spacing⌋, and an environment score is derived from the local density-weighted average of surrounding property grid voxels within a defined radius R. The fragment library is then queried for fragments whose interaction type matches the voxel environment, for example, hydrogen-bond donor fragments directed to regions of high acceptor potential. Fragment attachment vectors are aligned with the field gradient stored in the target pocket voxel, as managed by the attachment point manager, which evaluates hybridization (sp, sp^2^, sp^3^), aromaticity, and available valence. Throughout assembly, the validation engine enforces molecule constraints including maximum heavy atom count (10–50), molecular weight (150–700 Da), maximum LogP, and rotatable bond limits, ensuring drug-likeness is maintained at each growth step.

#### Multi-Objective Optimization and Scoring

Chemical space exploration is guided by a multi-objective scorer that produces a composite molecular score as a weighted linear combination: S_total_ = Σ(w_i_ × s_i_), where each component score s_i_ is multiplied by a user-configurable weight w_i_. The scoring components and their significance are summarized in Table 1. LigandForge is also equipped with present configurations that allow users to select tailored weighting schemes for specific target classes, such as kinases or GPCRs. These schemes automatically adjust the scoring weights within the multi objective scorer, prioritizing different combinations of pharmacophore alignment, synthetic accessibility, and drug-likeness to suit the unique structural and chemical requirements of the chosen protein family.

**Table 1.** Multi-objective scoring components used in the molecular score reward function.

| Metric | Significance |
| --- | --- |
| QED | Quantitative estimate of drug-likeness and lead-like properties. |
| Synthetic accessibility | Feasibility of the retrosynthetic route. |
| Pharmacophore alignment | Spatial complementarity to interaction hotspot locations. |
| Novelty and diversity | Structural uniqueness verified by enhanced diversity manager. |

To ensure that theoretically potent candidates remain practically synthesizable, a retrosynthetic penalty is applied to the final reward: S_ninal_ = S_total_ × (1 − D) × F, where D is the overall difficulty and F is the route feasibility from the retrosynthetic analyzer. Molecules exceeding predefined thresholds for total synthetic steps or overall difficulty receive a near-zero multiplier, effectively removing them from the viable optimization space. Optimization of the generated molecules is driven by one of three configurable algorithms; reinforcement learning, genetic algorithms, or a hybrid approach, and an optimization history object tracks population evolution, monitoring convergence metrics and population diversity to prevent stagnation in local optima.

#### Diversity Management and Retrosynthetic Feasibility

Scaffold diversity in the candidate output is maintained by the enhanced diversity manager, which employs DBSCAN^15^ clustering and Tanimoto similarity^16^ assessments on molecular fingerprints to ensure broad chemical space coverage. Final candidates undergo rigorous retrosynthetic analysis: the retrosynthetic analyzer decomposes target molecules into a series of synthetic steps, identifying required starting materials, reagents, and reaction conditions. Each proposed route is assigned a route feasibility score (F) and a synthetic complexity score, ensuring that output leads are both potent and chemically accessible.

#### Software Implementation

LigandForge is implemented in Python with a Streamlit-based front-end for interactive data exploration. All molecular manipulation, property calculation, and two-and three-dimensional structure rendering are performed using RDKit. Voxel grid construction and spatial analyses are carried out using NumPy and SciPy. Radar plots of multi-objective scores, property space distributions (MW vs. LogP), and optimization convergence trajectories are rendered using Plotly and Matplotlib. Runtime performance is monitored by psutil, and data consistency across all pipeline stages is enforced by a dedicated validation engine. The application is deployed as a web-based service on the Render cloud platform and is publicly accessible at: https://ligandforge.onrender.com.

### Validating LigandForge candidates against co-folding models

For the purpose of prospective validation, small-molecule ligands for dopamine receptor D2 (D2R), epidermal growth factor receptor (EGFR) and the N-terminal domain of STAT5b (STAT5b-NTD) were generated using LigandForge. De novo ligand design runs were conducted ten times for each receptor. At every run, the best ten molecules were collected based on the total score. This resulted in 100 generated molecules for each target, which were further evaluated with Boltz-2, a state-of-the-art co-folding model, for the prospective binding modes and affinities^8^. For D2R and EGFR, experimental structures from the Protein Data Bank (6CM4^11^ and 1M17^17^) were used as input for LigandForge. In the case of STAT5b-NTD a homology model based on the STAT3 N-terminal domain (PDB ID: 4ZIA) was used, which was described elsewhere^18^. The binding sites were selected based on the co-crystallized ligand present in the structure for D2R and EGFR. For STAT5b-NTD, in lack of a ligand, the average coordinates of the sidechain centroids for the residues of the handshake dimer interface were used as the grid center^19,20^. Binding site radii of 13.0 Å, 10.0 Å and 15.0 Å were used for D2R, EGFR and STAT5b-NTD, respectively. Kinase, GPCR and High quality configuration presets were used for D2R, EGFR and STAT5b-NTD, respectively. All interaction types were considered for molecule generation.

Ligand-protein complex structures and affinities were predicted using Boltz-2 for the selected LigandForge generated molecules with their respective protein targets. Input protein sequences for structure prediction were derived from the structures used as input for LigandForge. Pocket constraints were used for determining the binding of the ligand structure using the residues within 5.0 Å from the co-crystallized ligand for D2R and EGFR and using the handshake interface residues for STAT5b-NTD. Consistently with the convention of the software suite itself, affinity values for Boltz-2 predictions are reported as the logarithm of affinity, expressed in µM units, if not stated otherwise.

To assess the uniqueness of the molecules generated by LigandForge, we compiled sets of previously reported compounds with measurable activity against the receptors under investigation. In total, 1542 active compounds for D2R and 5956 active compounds for EGFR were retrieved from ChEMBL version 36^21^. As STAT5b-NTD is a considerably less explored target, only three active molecules with experimentally reported activities could be identified from our recent work^18^. We determined the most similar active for every generated molecule based on Tanimoto similarities^22^ of Morgan fingerprints (2048 bit, radius: 4).

## Conclusions

LigandForge offers a small-molecule drug discovery framework that integrates structure-guided design with synthetic feasibility in an accessible graphical user interface. By combining voxel-based property grids for high-resolution interaction hotspot characterization with a multi-objective scoring engine, the platform enables the systematic evolution of lead candidates that matches pharmacophoric requirements and drug-likeness. The platform’s Streamlit-based interface, featuring interactive NGL 3D visualization, further lowers the barrier to adoption, making sophisticated structure-based workflows accessible to researchers without deep computational expertise. Taken together, LigandForge establishes itself as a robust, end-to-end solution for accelerating hit identification and lead optimization across diverse therapeutic targets.

## AUTHOR INFORMATION

## DATA AVAILABILITY

Molecular structures and structure-guided design data, derived from the RCSB PDB and curated fragment libraries as described in this work, are made available through the LigandForge webserver (https://ligandforge.onrender.com). The platform supports interactive 3D visualization via an integrated NGL viewer and enables bulk download in multiple formats, including SDF and Excel. LigandForge facilitates the automated generation of lead candidates through voxel-based pocket analysis, multi-objective scoring, and retrosynthetic route prediction. The webserver is implemented in Python using the Streamlit framework, with RDKit for cheminformatics. The full source code is publicly available on GitHub (https://github.com/HTS-Oracle/LigandForge).

## ASSOCIATED CONTENT

The following files are available free of charge.

Two-dimensional representation of the displayed top-candidate molecules generated by LigandForge and the most similar active compound found in the literature (PDF).

SMILES strings, Total Score, Boltz Affinity, Max Similarity, Most Similar Active, and Target for all compounds (XLSX).

## Acknowledgments

This work was supported by the National Institutes on Aging under grant number RF1AG084635 (PI: Gabr). In addition, this work was also supported by the Servier-Beregi Scholarship and by the National Research Development and Innovation Office of Hungary [contracts FK146063 to D.B., NKKP 153477, 152137 and PharmaLab (RRF-2.3.1-21-2022-00015) to G.M.K.]. The work of D.B. was supported by the János Bolyai Research Scholarship of the Hungarian Academy of Sciences. We are grateful for the possibility to use the GenAI4Science service of the HUN-REN Cloud (see Héder et al. 2022; https://science-cloud.hu/) which helped us achieve the results published in this paper.

## Notes

The authors declare no competing financial interest.

## Insert Table of Contents artwork here

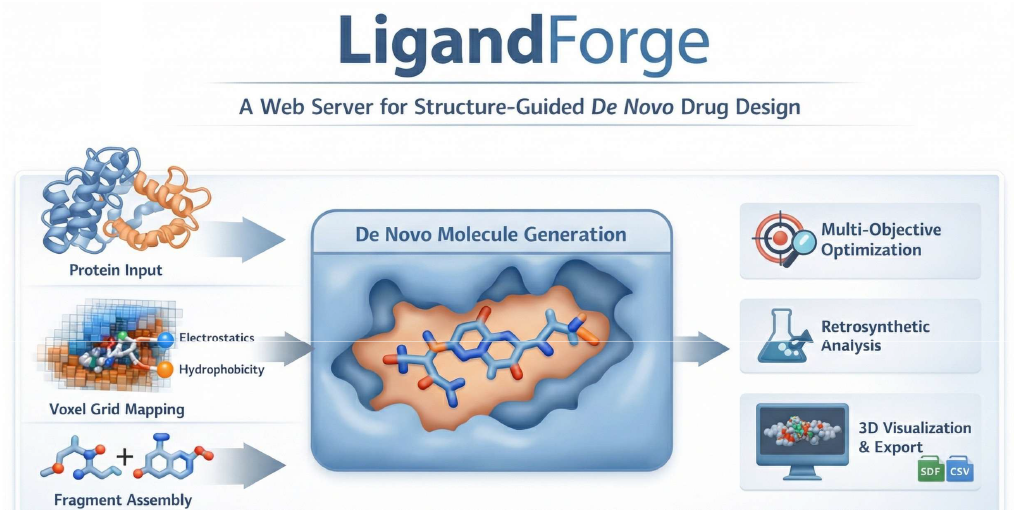

## Notes

### Competing Interest Statement

The authors have declared no competing interest.

