## Supporting Information for "LigandForge: A Web Server for Structure-Guided De Novo Drug Design"

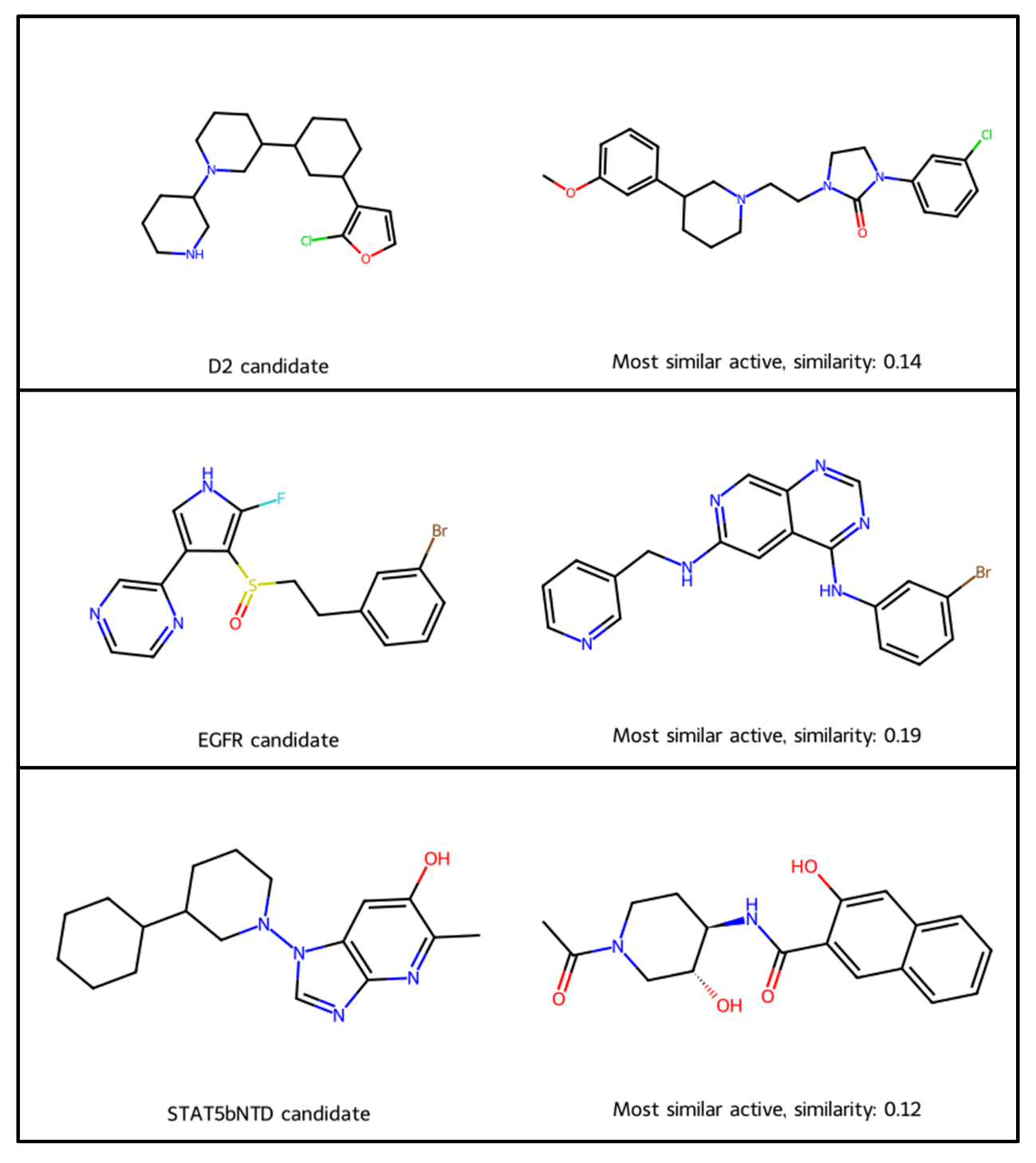

**Supplementary Figure S1.** Two-dimensional representation of the displayed top-candidate molecules generated by LigandForge for D2R, EGFR and STAT5b-NTD (left side). On the right side, the most similar active compound found in the literature is displayed (with their similarity value to the given candidate molecule). Similarity values are calculated based on the Tanimoto distance of Morgan fingerprints.
